# DeepRegFinder: Deep Learning-Based Regulatory Elements Finder

**DOI:** 10.1101/2021.04.27.441658

**Authors:** Aarthi Ramakrishnan, George Wangensteen, Sarah Kim, Eric J. Nestler, Li Shen

## Abstract

**Motivation:** Enhancers and promoters are important classes of DNA regulatory elements that control gene expression. Identifying them at the genomic scale is a critical and challenging task in bioinformatics. The most successful method so far is to train machine learning models on known enhancer and promoter sites and predict them at other genomic regions using ChIP-seq and related data.

**Results:** We have developed a highly customizable program called DeepRegFinder which automates data processing, model training and genome-wide prediction of enhancers and promoters using convolutional and recurrent neural networks. Our program further classifies the enhancers and promoters into active and poised states to facilitate downstream analysis. Based on mean average precision scores of different classes across multiple cell types, our method significantly outperforms the existing algorithms.

**Availability:** https://github.com/shenlab-sinai/DeepRegFinder

## 1. Introduction

DNA regulatory elements (DREs) are genomic regions that control gene expression through their interactions with chromatin- and DNA-binding regulatory protein. The major categories of DREs include promoters, enhancers, silencers and insulators, with promoters and enhancers being the ones most widely studied. Promoters are stretches of DNA that are proximal to the transcriptional start sites (TSSs) of genes and are approximately 0.1 to 1 Kb in length. Most promoters of protein coding-genes have already been identified. Enhancers on the other hand are DREs that are up to 1 Mb away from the TSSs of the genes that are being regulated (Pennacchio, et al., 2013). Enhancers are known to be cell-type-specific and closely involved in development and disease (Parker, et al., 2013; Spitz and Furlong, 2012). The ENCODE project (ENCODE Project Consortium, 2012) aims to identify all DREs – a large proportion of which being enhancers – across a large number of cell types (https://www.encodeproject.org/data/annotations/). Despite the progress that has been made, numerous enhancers have still not been characterized due to their dynamic nature. Both enhancers and promoters display distinctive chromatin modification patterns that are characterized by the enrichment and depletion of specific histone marks. This makes it possible to identify all enhancers and promoters in the genome, provided the binding profiles of different histone marks from ChIP-seq or related (e.g., CUT&RUN) data are available for each tissue of interest. Several machine learning methods (Kim, et al., 2016; Kleftogiannis, et al., 2015; Liu, et al., 2016; Lu, et al., 2015; Rajagopal, et al., 2013) have been developed which are trained on known enhancers and promoters to identify the unknown ones across the genome.

There are certain limitations to the existing methods for enhancer identification. First, for a machine learning task, the most time-consuming part often lies in data processing and transformation. However, most methods published to date have focused on model validation alone. The accompanying software for such methods is often hard to apply on large sequencing datasets obtained from diverse cell types. Second, most studies tend to single out enhancers as the positive class and group everything else into the negative class, including promoters. Because enhancers and promoters share common histone marks and ChIP-seq data contain intrinsic noise, it is much more challenging to distinguish enhancers from promoters than from background genomic regions. Therefore, model evaluation based on binary classification where the promoters are grouped together with the negative class will conceal the real performance in distinguishing different classes of DREs. Finally, because enhancer function varies considerably among different tissues, it is essential to identify enhancers based on data derived from the particular tissue of interest.

In this study, we present DeepRegFinder – a highly customizable computational pipeline that allows an investigator to process ChIP-seq data and build deep learning models to predict DREs using a user-friendly interface. Our program performs 5-way classification of active and poised enhancers (AEs and PEs, respectively) and promoters (ATs and PTs, respectively) plus background (Bgd) genomic regions. Our deep learning models use convolutional neural network (CNN) and recurrent neural network (RNN) (LeCun, et al., 2015), and we compare their performance with existing approaches.

## 2. Methods

DeepRegFinder is designed to be easy-to-use and highly customizable; requiring only the most commonly used file formats as input, such as BAM files (Li, et al., 2009) and text-based annotation files. It has three modules: preprocessing, training and prediction. Each module is a Python script that uses a text based YAML file as input, which specifies the input files and parameters. The preprocessing module takes inputs such as genomic annotation, peak lists and ChIP-seq alignments (i.e., BAM files) to build a training set for read coverage of known enhancers and promoters and a randomly selected set of background genomic regions. Each enhancer or promoter is further classified into active or poised states based on 2-way K-means clustering on GRO-seq (Core, et al., 2008) derived coverage. Each input sample represents a 2 Kb window that is separated into 20 equal bins (both window and bin size can be changed as parameters). The read coverage for multiple histone marks is calculated and normalized for each bin. The entire training set can therefore be encoded in a 3-D tensor of samples×histone marks×bins. The preprocessing module also calculates the read coverage for all bins that tile the whole genome to be used by the prediction module. The training module trains a deep learning model to perform 5-way classification using the read coverage as input. A user can specify the neural network structure and other training parameters such as learning rate, number of epochs and batch size. The prediction module runs the trained models on the genomic read coverage to perform classifications for each 2 Kb window in step of 100 bps. Each module can run independently, which allows a lot of flexibility in execution. For example, a user may train a model using public ENCODE data and later use the trained model to predict on her private data using the preprocessing and prediction modules—skipping the training module altogether.

The CNN and RNN structures are shown in Supplemental Fig. S1. Both structures use a 7×1 1-D convolution layer at the bottom as a feature extractor for histone modification patterns, which acts like a motif detector. For CNN, several 1-D convolution layers are stacked on top of one another, with two max pooling layers interspersed to reduce spatial resolution. A global average pooling layer is used, followed by a softmax layer for classification. For RNN, two long-short term memory networks (LSTMs) (Hochreiter and Schmidhuber, 1997) with the same size of 32 are stacked on top of the 1-D convolution layer, followed by a softmax layer for classification. All convolution layers are followed by batch normalization (BN) (Ioffe and Szegedy, 2015) and Rectified Linear Unit (ReLU) (Dahl, et al., 2013; Nair and Hinton, 2010). Both CNN and RNN use Adam (Kingma and Ba, 2014) as the optimizer.

## 3. Results and Discussion

The DeepRegFinder preprocessing module was used to create datasets for training, evaluation and genome-wide prediction on three different cell types – K562, H1 and GM12878 – downloaded from the ENCODE website (https://www.encodeproject.org/). Each dataset contained about a dozen different histone marks and for each of the four non-background classes, the number of samples varied between approximately 4,000 to 33,000 (Supplemental Table S1); for the background class, the number of samples was fixed at 100,000. Each dataset was split into train-validation-test sets based on a 60-20-20 proportion; class stratification was used to maintain the same proportions of the five classes across the train-validation-test sets. The validation set was used to choose best models and the test set was used for performance evaluation.

Unlike other studies, we did not use the AUC score for model evaluation because it can conceal or ignore problems with false positive determinations when the background class is much larger than the non-background classes. Instead, we used the mean average precision (mAP) of the four non-background classes as the evaluation metric. Both CNN and RNN were compared with two established methods – RFECS (Rajagopal, et al., 2013) and EP-DNN (Kim, et al., 2016). Table 1 lists the mAP scores of the four methods and Table S2 lists the average precisions for the four non-background classes separately. Both RNN and CNN show mAP scores at around 0.70, which are significantly better than that of RFECS and EP-DNN. Additionally, RNN’s performance was almost as good as that of CNN. Confusion matrix analysis (Supplemental Fig. S2) shows that most active enhancers and active promoters were correctly classified, while the poised enhancer and poised promoter classes had much lower accuracies. Supplemental Fig. S3 illustrates example screenshots for the predictions made by DeepRegFinder (CNN) in K562 cells, which show that the predicted enhancers and promoters exhibit characteristic binding patterns of particular histone marks.

**Table 1.** mAPs of the four comparing methods across three different cell types on the test set.

| mAP [95% CI] | K562 | H1 | GM12878 |
| --- | --- | --- | --- |
| **DeepRegFinder (CNN)** | 0.72 [0.71,0.73] | 0.69 [0.68,0.71] | 0.68 [0.67,0.69] |
| **DeepRegFinder (RNN)** | 0.72 [0.71,0.73] | 0.69 [0.67,0.70] | 0.67 [0.67,0.68] |
| **EP-DNN (Kim, et al., 2016)** | 0.62 [0.61,0.63] | 0.65 [0.64,0.66] | 0.63 [0.63,0.64] |
| **RFECS (Rajagopal, et al., 2013)** | 0.64 [0.63,0.65] | 0.67 [0.66,0.68] | 0.63 [0.63,0.64] |

One of the advantages of using a convolution layer as the feature extractor for the first layer is that the convolution layer’s weight parameters can be interpreted as filters for primitive patterns whose combinations are learned by upper layers for classification. Fig. 1 shows the clustering of the activations of the 32 filters of the first convolution layer (CNN trained on K562 cells) for the top 100 samples from each class and the associated weight parameters for different histone marks. Some filters are distinctly associated with specific classes. For example, filter 15 is associated with active promoters and contains peak detectors for H3K4me2/3, while filter 12 is associated with active enhancers and contains a peak detector for H3K27ac and is anti-correlated with H3K4me3.

**Figure 1.**
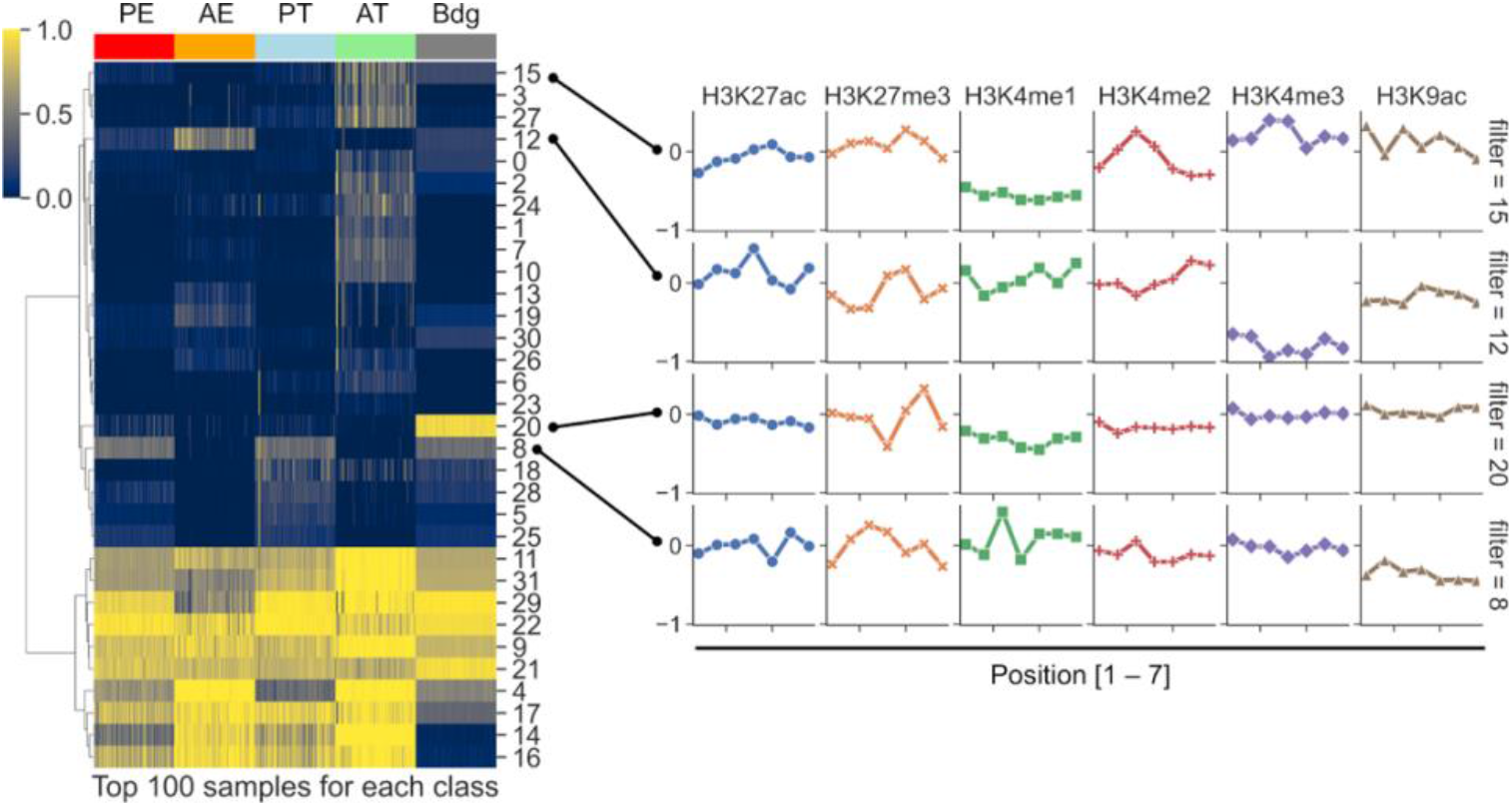
Clustering of the 32 filters of CNN trained on K562 cells based on the activations of the top 100 samples for each class. For the selected filters, their weight parameters for different histone marks are shown.

CNN and RNN in DeepRegFinder are both parameter efficient with only 26K and 12K weight parameters, respectively. In contrast, the Multi-Layer Perceptron (MLP) based EP-DNN has about 500K parameters. Running DeepRegFinder takes about 2-8 hr for preprocessing; 5 min for training; and 20 min for prediction for the entire human genome. Both training and prediction require only a modest GPU configuration.

## Acknowledgments

We would like to thank Dr. Weijia Zhang for providing the GPU server for conducting the research in this study.

