## Supplemental for "DeepRegFinder: Deep Learning-Based Regulatory Elements Finder"

**In total: 2 tables; 3 figures.**

*Table S1.* Basic statistics of the datasets used for training and evaluation. The background class size is 100,000 for all three cell types.

|  | PE | AE | PT | AT | Number of histone marks | List of histone marks |
| --- | --- | --- | --- | --- | --- | --- |
| H1 | 9508 | 4137 | 6701 | 11130 | 11 | H2AFZ, H3K27me3, H3K4me1, H3K4me3, H3K9ac, H4K20me1, H3K27ac, H3K36me3, H3K4me2, H3K79me2, H3K9me3 |
| K562 | 15172 | 18415 | 5403 | 14922 | 12 | H2AFZ, H3K27me3, H3K4me1, H3K4me3, H3K9ac, H3K9me3, H3K27ac, H3K36me3, H3K4me2, H3K79me2, H3K9me1, H4K20me1 |
| GM12878 | 25400 | 33461 | 11430 | 25665 | 11 | H2AFZ, H3K27me3, H3K4me1, H3K4me3, H3K9ac, H4K20me1, H3K27ac, H3K36me3, H3K4me2, H3K79me2, H3K9me3 |

*Table S2.* Average precisions with 95% confidence intervals for the four non-background classes for the three cell types.

|  | PE | AE | PT | AT |
| --- | --- | --- | --- | --- |
| Cell type = H1 | | | | |
| DeepRegFinder (CNN) | 0.629 [0.606, 0.653] | 0.530 [0.494, 0.567] | 0.693 [0.668, 0.716] | 0.922 [0.910, 0.933] |
| DeepRegFinder  (RNN) | 0.623 [0.599, 0.648] | 0.516 [0.480, 0.554] | 0.682 [0.657, 0.707] | 0.919 [0.907, 0.930] |
| EP-DNN (Kim, S.G. et al.) | 0.563 [0.538, 0.589] | 0.456 [0.421, 0.491] | 0.669 [0.645, 0.693] | 0.912 [0.899, 0.923] |
| RFECS (Rajagopal, N. et al.) | 0.552 [0.529, 0.578] | 0.464 [0.431, 0.503] | 0.650 [0.625, 0.676] | 0.907 [0.896, 0.918] |
| Cell type = K562 | | | | |
| DeepRegFinder (CNN) | 0.690 [0.672, 0.708] | 0.821 [0.808, 0.834] | 0.456 [0.425, 0.486] | 0.916 [0.907, 0.926] |
| DeepRegFinder  (RNN) | 0.690 [0.672, 0.708] | 0.821 [0.807, 0.834] | 0.451 [0.421, 0.481] | 0.907 [0.895, 0.919] |
| EP-DNN (Kim, S.G. et al.) | 0.562 [0.543, 0.582] | 0.737 [0.720, 0.754] | 0.294 [0.265, 0.326] | 0.881 [0.868, 0.894] |
| RFECS (Rajagopal, N. et al.) | 0.604 [0.586, 0.624] | 0.789 [0.775, 0.802] | 0.396 [0.366, 0.427] | 0.904 [0.894, 0.914] |
| Cell type = GM12878 | | | | |
| DeepRegFinder (CNN) | 0.624 [0.610, 0.639] | 0.824 [0.814, 0.833] | 0.399 [0.378, 0.420] | 0.865 [0.857, 0.874] |
| DeepRegFinder  (RNN) | 0.614 [0.600, 0.629] | 0.823 [0.813, 0.832] | 0.392 [0.371, 0.412] | 0.862 [0.853, 0.871] |
| EP-DNN (Kim, S.G. et al.) | 0.563 [0.549, 0.579] | 0.795 [0.785, 0.806] | 0.335 [0.315, 0.356] | 0.840 [0.830, 0.850] |
| RFECS (Rajagopal, N. et al.) | 0.551 [0.538, 0.566] | 0.794 [0.784, 0.804] | 0.330 [0.310, 0.351] | 0.837 [0.827, 0.846] |

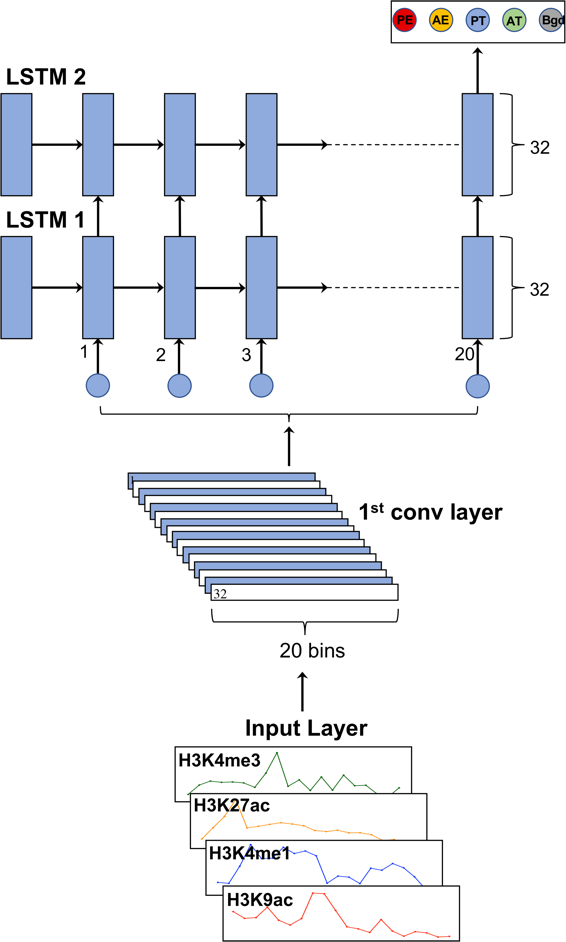

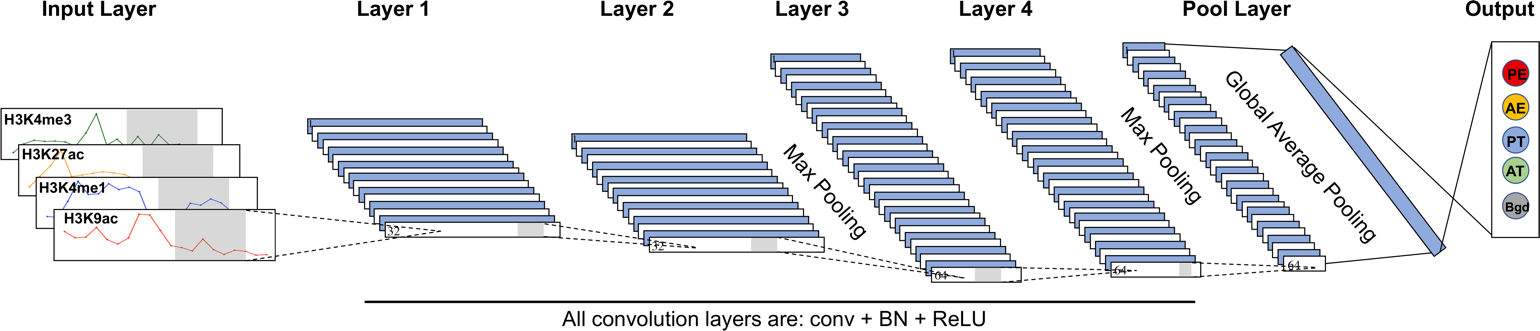

a

b

*Figure S1.* Network structures for a) CNN and b) RNN. For CNN, the max pooling layers reduce the spatial dimension by two fold and the global average pooling layer averages across all spatial positions.

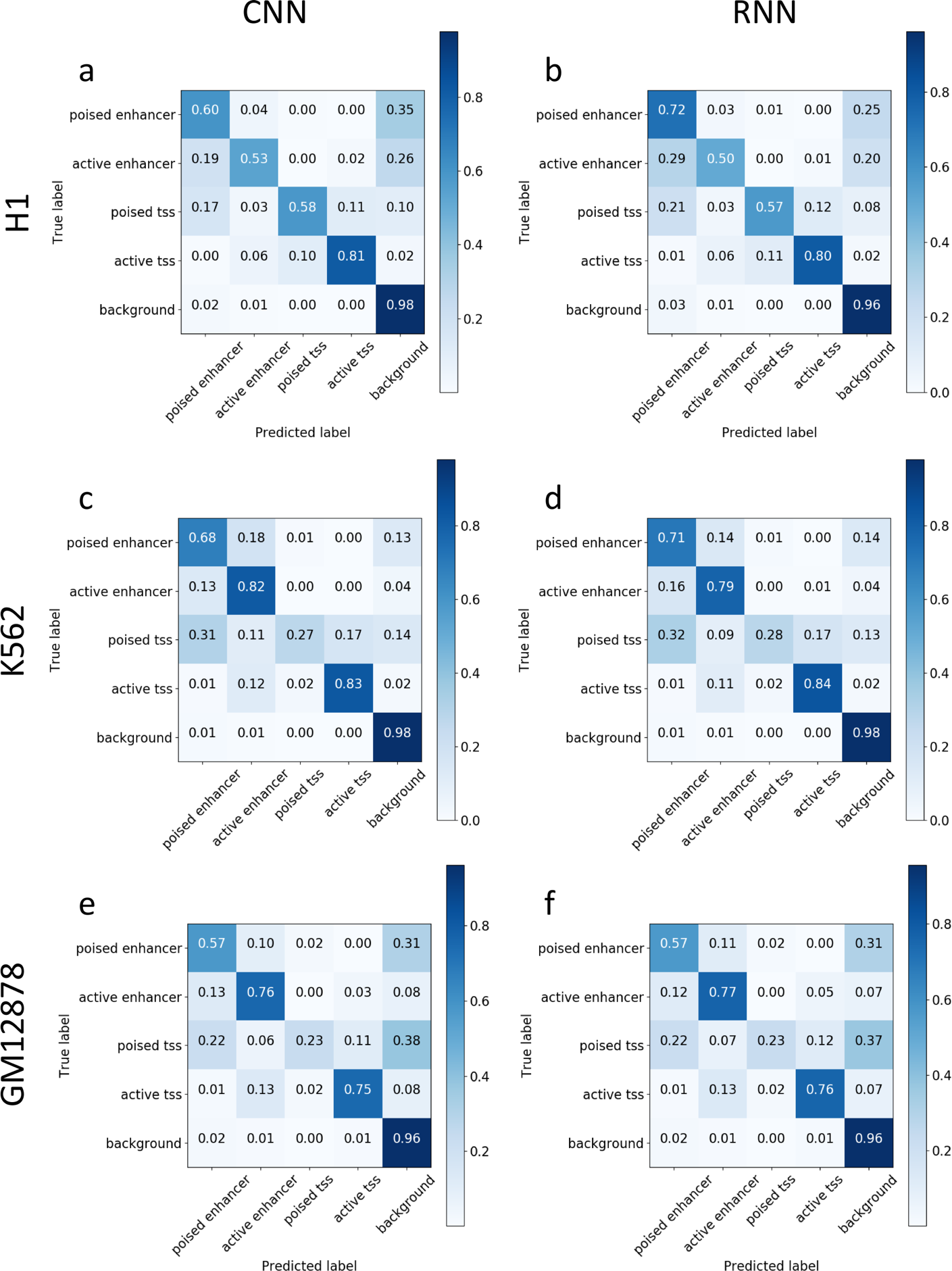

*Figure S2.* Confusion matrix analysis for the test set predictions in the three cell types – a) and b), c) and d), e) and f) for CNN and RNN, respectively.

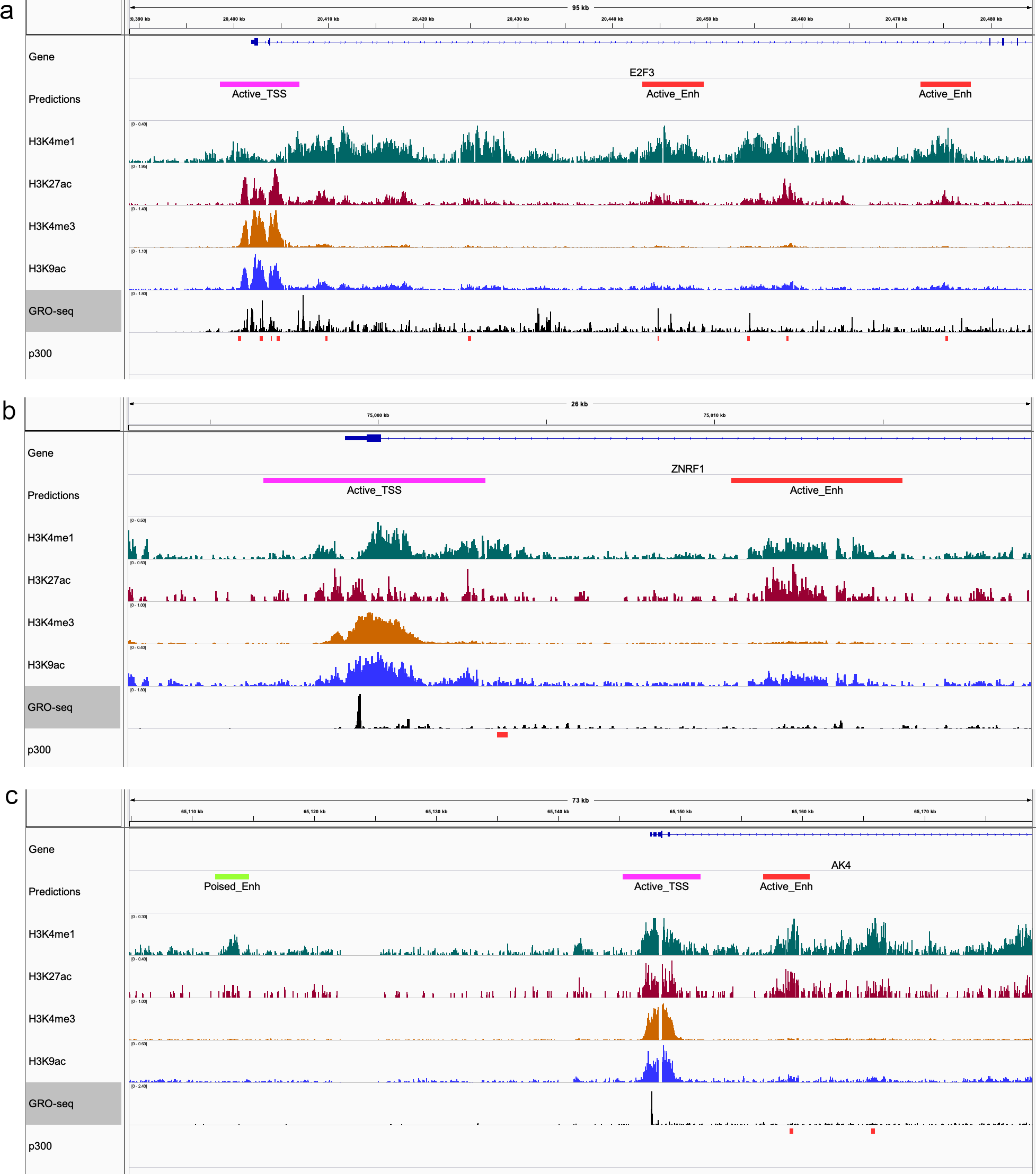

*Figure S3.* Example genome browser screenshots for predictions produced by DeepRegFinder (CNN) in K562 cells.
